# Ferrybots: Programmable Hydrogel Microbots for Rapid Magnetic Navigation and Modular Function

**DOI:** 10.64898/2026.11.11.763821

**Authors:** Vivek Rajasenan, Timothy Vernon, Ridhi Pradhan, Artem Goncharov, Hamed Hosseini-Toudeshki, Aiden Begole, Shun Ye, Thu Phung, Michael McShane, Aydogan Ozcan, Dino Di Carlo

## Abstract

Untethered microrobots could enable localized delivery, sensing, and intervention in confined biological and engineered environments. Magnetic actuation is especially attractive for such systems because it provides remote, fuel-free control, but practical magnetic microrobots remain difficult to manufacture and functionalize. High magnetic loading can interfere with polymerization and structural integrity, while integrating application-specific payloads directly into the robot body often requires the entire material formulation to be redesigned. Here, we introduce Ferrybots, programmable hydrogel microrobots that physically separate magnetic navigation from functional cargo. An automated droplet-on-demand microfluidic flow-lithography process confines high-concentration ferrofluid cores within photopolymerized poly (ethylene glycol) diacrylate (PEGDA) structures, enabling magnetic loading to be programmed through core number and spacing without uniformly dispersing magnetic particles throughout the hydrogel. The process achieved a minimum fabrication cycle time of 15 s. Ferrybots exhibited magnetic-dose-dependent translation and rotation, branched-channel steering, out-of-plane obstacle traversal, and rapid translation, with two-core robots following an external magnet at speeds up to the 9000 mm min⁻¹ limit of the actuation system. Differences in magnetic loading also enabled sequential selective navigation of distinct Ferrybot populations. Using PEGDA 575 as a common interface material, independently formulated hydrogel cargo modules could subsequently be fused to preformed Ferrybots while preserving navigation. Separately, cargo modules enabled composition-dependent molecular release and oxygen-responsive sensing through 1-mm-thick skin-mimicking optical phantoms representing a broad range of skin types. By decoupling locomotion from function, Ferrybots provide a common magnetic vehicle that can be combined with interchangeable delivery, sensing, or responsive-material modules, offering a general strategy for rapidly adapting microrobots to different applications.

## 1 Introduction

Microrobots capable of remotely controlled movement offer potential tools for localized delivery, sensing, environmental manipulation, and minimally invasive intervention^1–3^. However, their practical development remains challenging because propulsion, structural integrity, material compatibility, and scalable fabrication must often be optimized simultaneously within a single structure^4–6^. Although advances in semiconductor manufacturing, photolithography, three-dimensional printing, acoustic propulsion, and catalytic actuation have enabled increasingly sophisticated microrobotic systems, these approaches can require specialized actuation equipment, experience throughput limitations, or rely on operating conditions that complicate translation into biological environments^7–12^.

Magnetic actuation is particularly attractive because it enables fuel-free and remotely controlled movement using externally applied magnetic fields. Hard magnetic materials such as neodymium–iron–boron (NdFeB) can generate strong magnetic responses and support complex programmed deformations, but their use in biomedical microdevices raises material-safety and integration concerns^13–18^. Superparamagnetic iron oxide nanoparticles offer an alternative low-cost magnetic material and have been incorporated into microswimmers, grippers, and other polymeric microrobots^19–21^. However, uniformly dispersing high concentrations of magnetic nanoparticles throughout photopolymerizable matrices can promote aggregation, interfere with polymer-network formation, and compromise structural integrity^22–24^. As a result, many hydrogel-based magnetic microrobots balance magnetic loading against reliable polymerization and material performance.

Automated fabrication represents an additional challenge. Microfluidic manufacturing provides a route to automated production with precise control over particle geometry and composition^25,26^. However, most existing microfluidic approaches also incorporate magnetic material throughout the polymer body, coupling magnetic performance to the chemical and mechanical requirements of the surrounding matrix. Furthermore, delivery or sensing functions are often integrated directly into the magnetic body, requiring the complete robot formulation to be redesigned for each application^27^. A modular strategy that spatially confines magnetic material while allowing independently optimized functional hydrogel cargo to be attached to a common vehicle could therefore simplify the development of multifunctional microrobotic systems.

Here, we introduce Ferrybots, superparamagnetic nanoparticle (Fe_3_O_4_) ferrofluid-core hydrogel microbots fabricated using a droplet-on-demand microfluidic flow-lithography platform (Fig. 1). Rather than uniformly dispersing magnetic nanoparticles throughout the polymer matrix, the approach spatially encapsulates programmable numbers of high magnetic-content liquid ferrofluid cores within photopolymerized PEGDA microstructures. This architecture enables control over magnetic loading while preserving a hydrogel exterior that can support subsequent attachment of independently formulated cargo. We first establish controlled ferrofluid core generation, hydrodynamic focusing, and automated Ferrybot fabrication. We then characterize magnetic-dose-dependent translation, rotation, rapid movement, selective actuation, branched-channel steering, and 3-dimensional obstacle traversal.

**Fig. 1.**
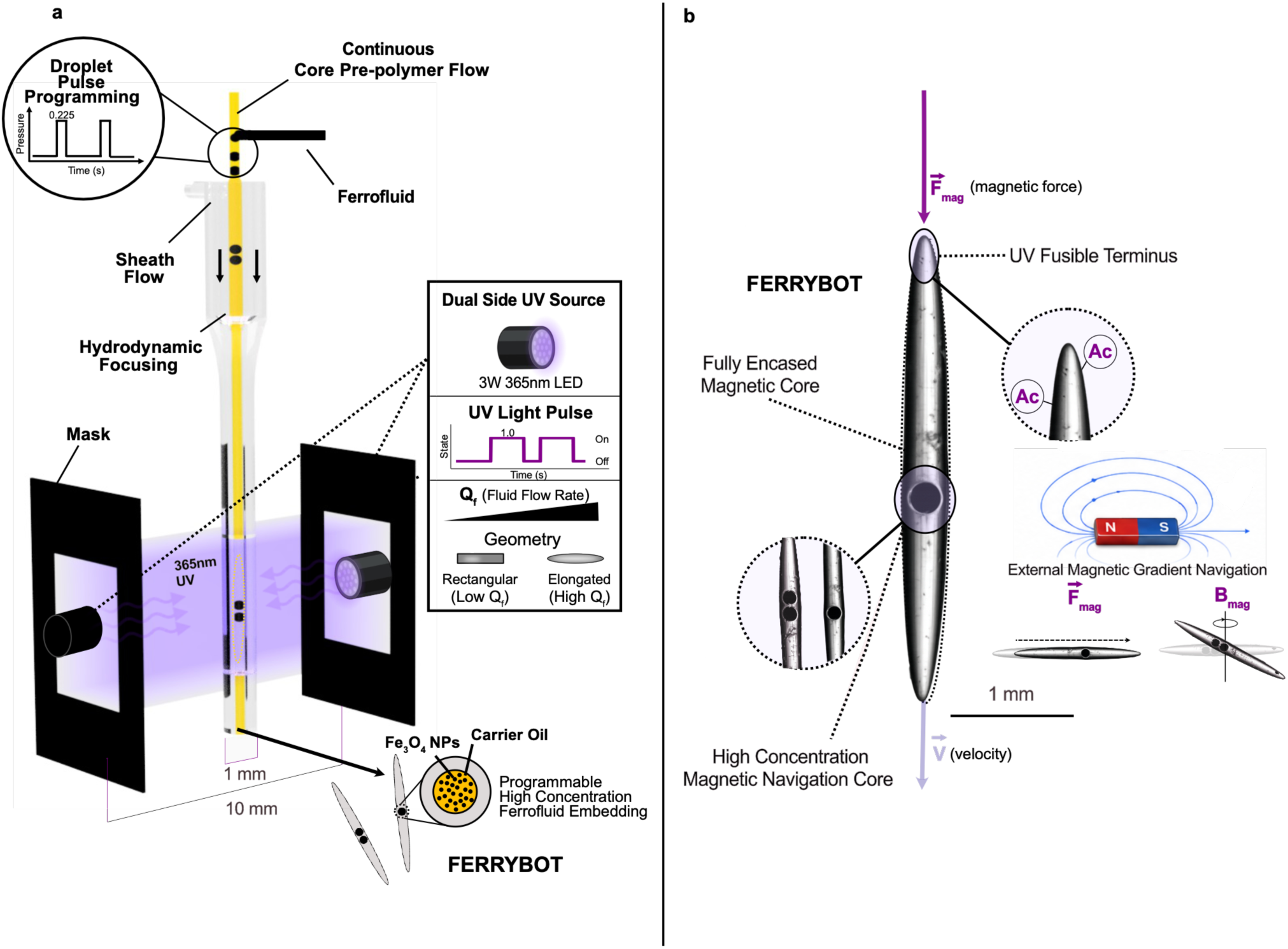
Overview of fabrication and magnetic actuation of hydrogel Ferrybots. (a) Droplet-on demand system produces elongated hydrogel microstructures with high dose magnetic navigation cores via liquid encapsulation and spatially controlled UV photolithography. (b) Embedded magnetic navigation cores in conjunction with external magnetic field gradients allow programmable remote magnetic actuation for translation and rotational movement. Definitions: Ac – acrylate group; F_mag_ – magnetic force; B_mag_ – magnetic field strength

To extend the platform beyond locomotion, we developed a secondary fabrication strategy for attaching PEGDA 575 hydrogel cargo to preformed Ferrybots while retaining magnetic mobility and channel-navigation capability. Rhodamine B was selected as a model small molecule to evaluate polymer-composition-dependent release, whereas oxygen-responsive PdBP nanoparticles were selected to demonstrate a fundamentally different phosphorescence lifetime-based sensing function. Oxygen was chosen as a physiologically relevant model analyte because local oxygenation reflects tissue perfusion, metabolism, and hypoxic state associated with various disease processes^28–30^. The oxygen-responsive modules were additionally evaluated through 1-mm-thick skin-mimicking optical phantoms to determine whether analyte-dependent lifetime contrast could be recovered through absorbing and scattering media. Together, these studies establish Ferrybots as a modular magnetic hydrogel microbot platform that combines programmable magnetic loading and rapid navigation with an attachment strategy for interchangeable delivery- and sensing-related hydrogel cargo.

## 2 Materials and methods

### 2.1 Materials and reagents

The Ferrybots were fabricated using neat poly (ethylene glycol) diacrylate (PEGDA 250, *M*n = 250, Sigma-Aldrich) containing 3% (v/v) 2-hydroxy-2-methylpropiophenone photoinitiator (PI,HMPP; Darocur 1173, 164.2 g mol⁻¹, Sigma-Aldrich). The ferrofluid used in the embeddings was a low-viscosity oil-based ferrofluid (Ferrotec EFH1). The porous hydrogel cargo connected to the Ferrybot was fabricated using PEGDA 575 (*M*n = 575, Sigma-Aldrich) diluted to final concentrations of 50–95% (v/v) with 3% (v/v) HMPP in Dulbecco’s Phosphate-Buffered Saline (DPBS, Gibco). All solutions were prepared in 5 mL Eppendorf tubes that were covered in aluminum foil and stored at 4 °C until use to prevent polymerization in ambient light.

### 2.2 Fabrication of 3D-printed microfluidic components

Flow-focusing nozzles and auxiliary microfluidic components were designed in Fusion 360 and exported as STL files. Flow-focusing nozzle designs were sliced in CHITUBOX using a 50 µm layer height and printed in Standard+ resin (Anycubic) using a masked stereolithography (MSLA) printer (Phrozen Sonic Mini 8K), with the nozzle tips oriented away from the build plate. The high-resistance, low-capacitance fluidic element described in Section 2.4 was fabricated by stereolithography using Standard+ resin and a 50 µm layer height, followed by press-cure bonding as previously described.^31^

Following printing, all microfluidic components were flushed three times with 100% ethanol and dried with compressed air to remove residual uncured resin. Channel patency was confirmed by visually verifying ethanol flow through the devices. The components were then post-cured for 15 min in a resin-curing chamber (SUNLU RC-2). STL files for the microfluidic device geometries are provided in the Supplementary Material (Schematic S1).

### 2.3 Simulation

To predict the focusing conditions and determine the acceptable operating space, single phase laminar flow simulations were conducted using COMSOL Multiphysics software (COMSOL Multiphysics, Burlington, MA). The geometry used for simulation was the inner nozzle channel drafted in the supplemental nozzle STL file (Schematic S1). The inlet conditions were set as constant flow rate boundary conditions. Sweeps of varying flow rate ratios of the sheath and core flows were conducted to calculate the predicted core flow diameter at the outlet. Single phase simulations were sufficient as core and sheath polymers are both comprised of ≥97% (v/v) PEGDA 250. Fluid viscosity (13.86 cP) and density (1.11 g/mL) were used for simulation^32^.

### 2.4 High-resistance, low-capacitance fluidic element

A high-resistance, low-capacitance fluidic element was incorporated upstream of the ferrofluid junction to enable rapid and reproducible conversion of short pressure pulses into discrete changes in ferrofluid flow. High hydraulic resistance reduces unintended ferrofluid flow arising from small pressure fluctuations, whereas low fluidic capacitance minimizes response delays associated with deformation of the device or connecting channels. Together, these characteristics enable discrete ferrofluid droplets to be generated on demand while minimizing trailing flow between pressure pulses.

The element contained 150 µm × 150 µm channels and was fabricated as described in Section 2.2. The rigid, press-cure-bonded structure was selected to minimize channel deformation and leakage under the elevated pressures associated with the high hydraulic resistance. This combination of high resistance and low compliance provided the rapid fluidic response required for programmable ferrofluid droplet generation.

### 2.5 Magnetic field motion system

To enable controlled translation and steering of the Ferrybots, we developed a three-axis motion system that moved a permanent magnet relative to a fixed microfluidic device. This configuration provided programmable control over the position and speed of the magnetic field source and supported both predefined trajectories and real-time manual navigation. The magnetic field was generated using a neodymium N52 permanent magnet mounted on a gantry-style translation system. NEMA 17 stepper motors controlled the magnet position along the x-, y-, and z-axes. The motors were driven by an SKR Mini E3 control board, which provided independent or coordinated motion along all three axes. The system was connected to a computer running custom Python software that accepted user-defined positions and translation speeds. These commands were converted to G-code and transmitted to the controller board through serial communication in real time^33^. Ferrybots could be steered using either keyboard inputs or a console controller. During operation, the microfluidic vasculature devices were secured in a 3D-printed holder at fixed coordinates within the motion system.

### 2.6 Ferrybot fabrication

The Ferrybot fabrication system combined four coordinated components: coaxial core and sheath prepolymer flows, pressure-pulsed ferrofluid droplet generation, temporally controlled UV photopolymerization, and downstream collection. Together, these components enabled ferrofluid droplets to be positioned within a focused photopolymerizable stream and encapsulated within individual hydrogel bodies. The core and sheath streams were both composed primarily of PEGDA 250 to provide similar fluid properties and support stable hydrodynamic focusing with minimal interfacial disturbance. The core flow stream contained PEGDA 250 with 3% (v/v) HMPP photoinitiator, whereas the sheath stream contained PEGDA 250 without photoinitiator and was therefore not intended to participate substantially in photopolymerization. Both streams were delivered using syringe pumps (Harvard Apparatus). The core and sheath flow rates were 15 and 35 µL min⁻¹, respectively, corresponding to a sheath-to-core flow-rate ratio of 2.33 and a total flow rate of 50 µL min⁻¹.

Ferrofluid droplets were introduced into the core stream using a pressure-pulse system composed of a two-way solenoid valve, a precision pressure regulator (Bellofram Marsh), and a compressed-air source (California Air Tools). Ferrofluid was loaded into a six-port stopcock manifold connected to the pressure source through the solenoid valve. After loading through one of the Luer-lock ports, the manifold was closed so that regulated air pressure directed the ferrofluid through the high-resistance fluidic element and into the droplet-generation junction. Solenoid actuation was synchronized with the fabrication control system. A 225 ms pressure pulse generated one ferrofluid droplet; correspondingly, a continuous 450 ms pulse generated two droplets under the selected operating conditions.

Photopolymerization was performed using two 3 W, 365 nm UV LEDs positioned on opposite sides of the nozzle outlet with 3D-printed holders. The LEDs were connected to an Arduino-based control system that synchronized UV exposure with the fluid velocity and the position of the ferrofluid droplets within the exposure region. The minimum UV exposure time used to fabricate structurally intact Ferrybots was 1 s.

Fabrication was coordinated using custom control software written in Python with the Tkinter graphical user interface library. The interface enabled users to specify the ferrofluid pressure-pulse duration, UV exposure duration, and delay between droplet generation and UV exposure. A live video feed was incorporated to monitor ferrofluid droplet formation, flow focusing, and positioning within the polymerization region during operation.

Following polymerization, Ferrybots were collected directly into 100% ethanol to remove residual unreacted monomer. The minimum total fabrication cycle time, including ferrofluid injection, droplet positioning, and UV polymerization, was 15 s per Ferrybot.

### 2.7 Imaging and analysis

Imaging of Ferrybots and their directed motion was conducted using a 5-megapixel microscope camera (Hayear, HY-500B). Video was taken at 30 frames per second (FPS). Ferrybots were tracked using the OpenCV package in Python, utilizing a background subtraction and image thresholding algorithm. Thresholding was adjusted until Ferrybots were clearly visible. Once isolated, a contouring step detected the center of the bot, which was tracked in each frame. The tracking data were then converted to physical units using a pixel-to-length calibration determined at the known camera working distance. Upon distance conversion, velocity and acceleration metrics were calculated relative to changes in Ferrybot distance with respect to time.

### 2.8 Polydimethylsiloxane (PDMS) mold fabrication for modular cargo integration

A reusable PDMS mold was fabricated to define the geometry of the modular hydrogel cargo added to preformed Ferrybots. The mold contained a 200 µm cylindrical channel that could be filled with a functional PEGDA 575 precursor solution while positioning the end of a Ferrybot within the channel. Subsequent photopolymerization formed the cargo module directly onto the Ferrybot, providing a controlled and reproducible interface between the magnetic vehicle and the independently formulated hydrogel cargo.

A master mold was designed in Fusion 360 and fabricated from poly (lactic acid) (PLA) by fused deposition modeling. The PLA structure was used to cast the PDMS mold and was selected because it permitted straightforward release of the cured PDMS^34^. The master contained a 108 mm × 16 mm cavity, producing a final PDMS mold thickness of 3.5 mm.

To form the cargo-molding channel, a 200 µm-diameter nichrome wire (Master Wire Supply) was inserted through aligned openings on opposite sides of the PLA master. The wire was pulled taut and secured to maintain a straight channel across the mold. PDMS base and curing agent were mixed at a 10:1 ratio and poured into the master around the wire. The assembly was cured at 65 °C for 3 h. After curing, the PDMS was removed from the PLA master and the nichrome wire was withdrawn, leaving a 200 µm cylindrical channel for positioning Ferrybots and molding the attached hydrogel cargo.

### 2.9 Small-molecule-eluting Ferrybot cargo

#### 2.9.1 Cargo fusion

Cargo-fused Ferrybots were fabricated by positioning previously formed Ferrybots within a PDMS mold (Section 2.8) containing PEGDA 575 with 3% (v/v) HMPP. The mold was then placed approximately 10 mm from UV source and exposed to a 3 W, 365 nm LED to polymerize the hydrogel and produce the fused cargo region on the Ferrybot.

#### 2.9.2 Rhodamine B cumulative release analysis

Rhodamine B cargo release from the PEGDA 575 hydrogel body was evaluated using a microplate reader (Infinite M Plex; Tecan). Cargo sections measuring 0.2 mm in diameter and 6.35 cm in length were fabricated by UV polymerization at 156 mJ/cm^2^ using PEGDA 575 concentrations of 50%, 80%, and 95% (v/v), each containing 0.04 mg/mL rhodamine B and 3% (v/v) HMPP. The 6.35 cm length approximated the inner circumference of the well, so each fiber rested against the well wall and the diffusion path to the measurement region at the center of the well was consistent across samples. The cargo sections were placed in identical volumes of 1× phosphate-buffered saline (PBS; Gibco) in 12-well plates and maintained at 37 °C. Each hydrogel cargo was positioned along the edge of its well to minimize interference with absorbance measurements collected from the center of the well. Absorbance at 554 nm was measured every 15 min for 48 h using three independent replicates per condition (n = 3). Cumulative release was expressed as the background-corrected absorbance at each time point normalized to the plateau absorbance measured for the corresponding PEGDA condition.

### 2.10 Oxygen detection cargo

#### 2.10.1 Oxygen Concentration Control

Controlled oxygen conditions spanning 0–21% O₂ were prepared in DPBS for characterization of the oxygen-sensing cargo. Here, oxygen percentage refers to the volumetric oxygen fraction of the gas phase with which the buffer was equilibrated. Air-equilibrated DPBS was used as the 21% O₂ condition, corresponding to approximately 257 µM dissolved oxygen at 25 °C and 210 µM at 37 °C.

The oxygen concentrations of 0–257 µM (0–21%) were generated by bubbling nitrogen gas through the buffer at controlled rates to partially displace oxygen. Dissolved oxygen concentrations were measured at the experimental temperature using a calibrated dissolved-oxygen probe (UNISENSE), with nitrogen-bubbled conditions characterized as described in the supplementary material of prior work^35^. All conditions were tested using three independent replicates (n = 3).

#### 2.10.2 Oxygen-sensing cargo fabrication

Oxygen-sensing cargo was fabricated by embedding oxygen-sensitive ethyl cellulose nanoparticles loaded with palladium (II) meso-tetra(4-carboxyphenyl) tetrabenzoporphyrin (PdBP-ECNPs) into 300 µm PEGDA 575 filled silicone tubing at a final concentration of 4 mg/mL with 3% (v/v) HMPP and UV cured at 365 nm with 156 mJ/cm^2^ (OmniCure). This phosphor was selected due to its long, red-shifted excitation and emission spectra for tissue penetration and long phosphorescence lifetime (∼230 µs) that decreases in proportion to oxygen concentration through collisional quenching, making it an intrinsic oxygen probe^36,37^.

#### 2.10.3 Flow chamber fabrication and perfusion for oxygen sensor testing

Flow chambers were fabricated from laser-cut acrylic slabs (75 × 25 × 1 mm). A rectangular fluidic channel (0.5 mm height) was cut at the center of one slab. The chamber was sealed using double-sided optically clear adhesive (3M) cut to match the channel geometry and enclosed with a second solid acrylic slab (75 × 25 × 1 mm) and transparency sheet (75 × 25 × 0.1 mm) to complete the flow cell. Inlet and outlet ports were connected to silicone tubing to enable controlled leak-free perfusion of test solutions through the chamber.

#### 2.10.4 Oxygen analyte characterization

The oxygen response was evaluated using the linear Stern–Volmer equation:

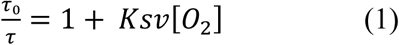

where *τ*₀ is the lifetime in the absence of oxygen, *τ* is the lifetime at a specific oxygen concentration, and the slope of the line, K_SV_, is the Stern–Volmer constant^38^. In the absence of oxygen, the phosphorescent lifetime is theoretically at a maximum value due to the lack of collisional quenching of the excited triplet state by molecular oxygen. Therefore, K_SV_ serves as a quantitative measure of the phosphor’s sensitivity to oxygen and dictates lifetime changes per unit oxygen concentration.

The phosphorescent lifetime was recorded using custom time-domain optical interrogation system with 630 nm excitation and 810 nm emission in a previously described flow-through system^39^. The samples (n = 3) were placed in a previously described custom-built flow cell consisting of an acrylic sheet that holds samples inside and optical readers attached from outside^39^. Lifetime was fit using an exponential decay function. Stern–Volmer fits were performed per condition across three replicates reporting mean and standard deviation. Lifetimes were collected once every 10 s with average lifetime determined via exponential fitting of the phosphorescence lifetime decay function as described in prior works^35^.

#### 2.10.5 Oxygen measurements through skin phantoms

To evaluate whether oxygen-dependent phosphorescence lifetime signals could be recovered through tissue-like optical attenuation, oxygen-sensing hydrogel cargo was measured through skin-mimicking phantoms. This experiment was designed to distinguish the effect of optical absorption and scattering on signal recovery from changes in the oxygen environment surrounding the sensor. Skin phantoms with a thickness of 1 mm were provided by collaborators and designed to reproduce the reduced scattering coefficient, μ′ₛ, and absorption coefficient, μₐ, associated with Fitzpatrick skin types I–VI at 630 nm excitation, based on previously reported optical-property targets^40^. The phantoms were grouped as Fitzpatrick types I–II, III–IV, and V–VI to represent low, intermediate, and high optical attenuation conditions. For each measurement, a skin phantom was placed directly atop the flow chamber containing the oxygen-sensing hydrogel cargo, creating a controlled optical barrier between the sensor and the detection system. Phosphorescence lifetime and intensity were measured during continuous perfusion of defined oxygen conditions (0% and 10% O₂), as described above. Across phantom groups, the oxygen-defined reagent, flow rate, temperature, hydrogel cargo, and sensor-placement geometry were held constant. The phantom therefore altered only the optical path, while the oxygen concentration surrounding the sensing cargo remained unchanged for a given condition. Review previous works for visual reference to the flow-chamber and phantom-placement geometry described ^35,37^. Three cargo replicates were measured for each condition (n = 3). A single phantom specimen was used for each Fitzpatrick-type group across experiments.

## 3 Results

The Ferrybot platform was designed around a compartmentalized fabrication strategy that separates magnetic loading from the surrounding hydrogel structure. Rather than dispersing magnetic nanoparticles throughout the polymer matrix, discrete ferrofluid droplets were generated, positioned within a focused PEGDA prepolymer stream, and subsequently encapsulated by spatially controlled photopolymerization. This approach was intended to enable programmable magnetic content while preserving a continuous hydrogel body suitable for navigation and subsequent cargo attachment.

We therefore first established control over ferrofluid droplet generation, number, and spacing using a pressure-driven droplet-on-demand system. We then integrated this process with hydrodynamic focusing and UV polymerization to fabricate Ferrybots with defined magnetic loading. After establishing fabrication, we evaluated magnetic actuation, navigation, selective control of distinct Ferrybot populations, and integration of modular functional cargo.

### 3.1 Ferrofluidic droplet-on-demand system for controlled magnetic loading

Precise control over ferrofluid droplet formation was required to define the magnetic loading of each Ferrybot. We therefore developed a droplet-on-demand (DoD) system that used short pressure pulses to generate discrete ferrofluid droplets within the flowing PEGDA 250 prepolymer core stream. Determining acceptable flow conditions to control ferrofluid droplet generation within a prepolymer sheath flow was key to successful implementation of our approach, given the need for precise fluid manipulation. As shown in Fig. 2a, we started with a DoD system consisting of a T-junction with a 0.3 mm diameter channel, whose capillary pressure barrier helps suppress trailing ferrofluid droplets caused by pressure fluctuations. With this design a droplet will only generate when the pressure difference between the dispersed and continuous phases exceeds the Laplace pressure, 2γ/r. Here γ is the interfacial tension between the ferrofluid and PEGDA 250, measured to be ∼4.9 mN/m and r is the radius of curvature, in this case half the channel diameter (0.15 mm), giving 2γ/r ≈ 65 Pa. The pressure control system (Section 2.6) is connected to the T-junction via a high resistance low capacitance element (Fig. 2a; Schematic S1). The increased resistance for the dispersed phase allows for source pressure reduction and more stability in the ferrofluid droplet formation process^41,42^. Additionally, the rigidity minimizes the capacitance, affecting the formation speed since flow development time is proportional to ∼RC. Here R is the fluidic resistance, and C is the fluidic capacitance, and therefore having a low capacitance rigid system enables a reasonably high dispersed phase channel resistance needed for a stable ferrofluid droplet formation process while still enabling reasonably fast response times necessary for programming the flow. To achieve a stable continuous phase flow, we used a syringe pump to maintain the flow rates of the co-flowing prepolymer phases.

**Fig. 2.**
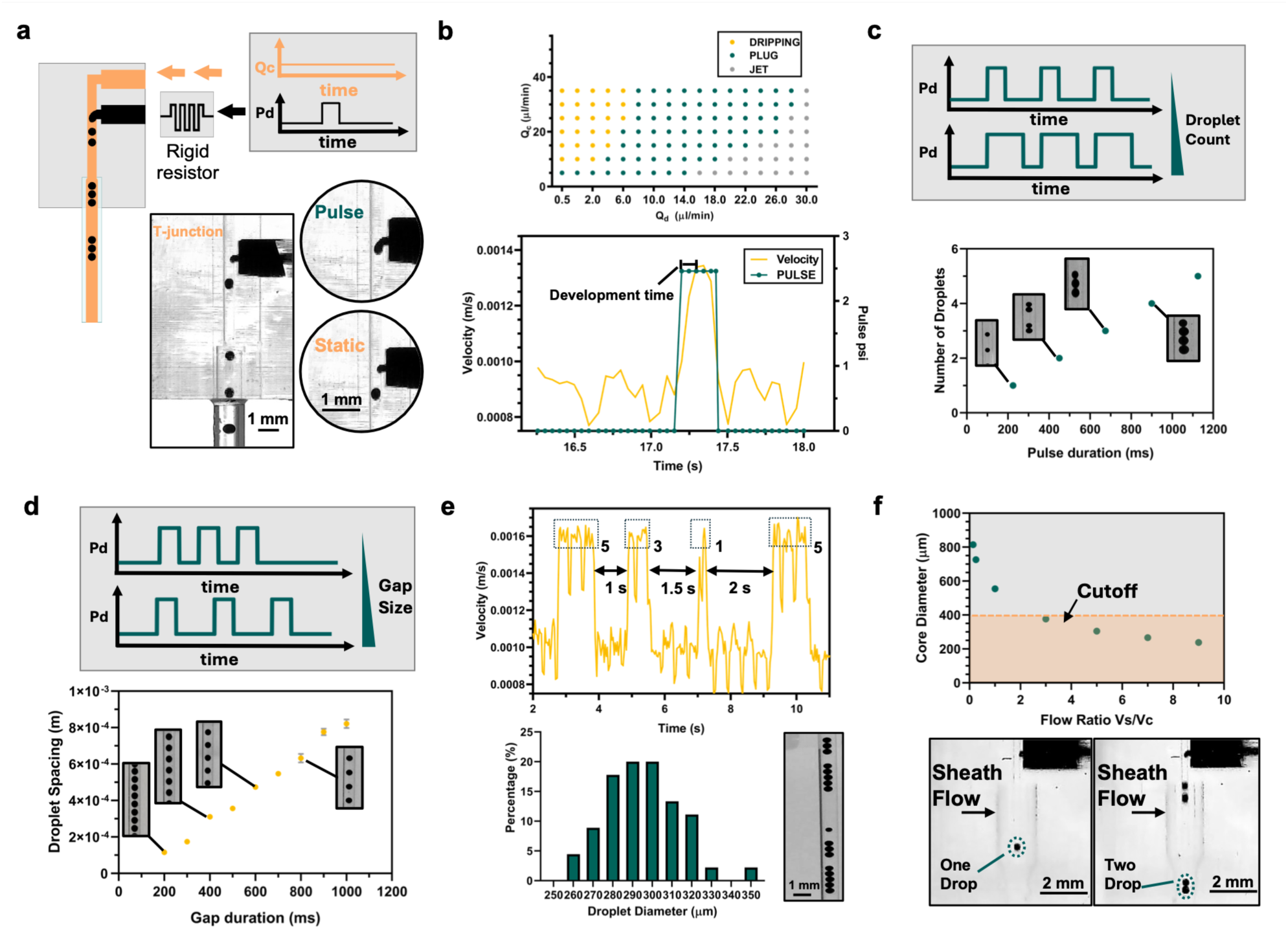
Flow characterization for the ferrofluid droplet-on-demand process. (a) Schematic and micrographs of the T-junction for droplet-on-demand (DoD) generation of ferrofluid droplets. High resistance rigid chip added for minimizing response time. (b) (top) Flow map for ferrofluid droplet generation demonstrating dripping, plug and jet regimes. (bottom) Velocity profile of DoD system overlapped with pressure pulse generated for ferrofluid droplet formation. (c) (top) Schematic of pressure pulse over time for increasing droplet number. (bottom) Linear increase in number of ferrofluid droplets for pressure pulse duration. (d) (top) Schematic of pressure pulse over time for increasing gap size between ferrofluid droplets. (bottom) Linear increase in gap size for increasing gap between pulses. (e) (top) ferrofluid droplet velocity across the formation of ‘5-3-1’ pattern. (Bottom) ferrofluid droplet diameter distribution and ferrofluid droplet pattern micrograph. (f) (top) Simulation of core-flow diameter for increasing flow rate ratios. (bottom) One and two droplet formations with full setup using sheath flow. Error bars denote standard deviation (n = 3). Scale bar = 2mm

Next, we sought to characterize the flow to determine the operating regime for the DoD system. Using a flow map of the continuous and dispersed phases flow rates (Fig. 2b (top)) we identified three formation modes for the ferrofluid droplets: ‘jetting’, ‘dripping’ and ‘plug’ flow. Since DoD requires active control, the system is unlikely to operate consistently in the jetting regime, due to the lack of a stable formation mode. The plug mode, while controllable, resulted in large ferrofluid droplets that span the channel width and would be difficult to fully encapsulate with the co-flowing sheathing polymers. Additionally, the change in pulse duration does not lead to a change in the number of droplets but rather the length of the plug formed. Based on our findings, we identified the dripping regime for future operation.

We were able to create a ferrofluidic DoD system with a development time of about 100 ms (Fig. 2b (bottom)), which was sufficient for programmability. With this capability at hand, we were able to program the number of droplets in a train following an expected linear trend with pulse width. Using an initial pulse duration of less than 225 ms resulted in either no droplet or an incomplete formation. Since a pulse width of 225 ms produced a single droplet on demand we expected to see an increase in droplet numbers for every 225 ms increment to the pulse width, which is reflected in experimental observations shown in Fig. 2c. The spacing between the droplets could also be controlled in a linear manner as shown in (Fig. 2d) by controlling the pressure pulse width and the width between the pulses. To demonstrate programmability, we produced a complex “5-3-1” droplet pattern with increasing gap spacings of 1, 1.5, and 2 s shown in Fig. 2e. The velocity and image data of the tracked droplets showed that the desired pulse pattern was generated. The generation of a pressure pulse resulted in a jump in the overall flow velocity due to the additional flow infusion from the ferrofluid dispersed phase. As expected, the resulting droplet pattern matched the design accurately with the creation of a 5-3-1 droplet train with increasing spacings between the droplet trains. Analysis of the droplet sizes across trials demonstrated a highly consistent process with an average droplet diameter of 294.8 µm and a coefficient of variation (CV) of 6.24% operating in the dripping regime shown in Fig. 2b. Having demonstrated a controllable ferrofluidic droplet-on-demand system, we next sought to incorporate the prepolymer sheath flow into the system.

The entire 3D-printed structure consisted of a T-junction combined with a prepolymer sheath flow that wraps around the T-junction to form an axisymmetric flow. Starting with CFD simulations (Fig. 2f), it was possible to deduce flow rate ratios of the core and the sheath flow under which the system would operate in ‘focusing mode.’ Under this mode the core flow maintains or reduces in diameter downstream relative to the nozzle diameter, a requirement for hydrodynamic focusing to have an effect since an increase in core-flow diameter destabilizes the droplet trains. It can be seen in Fig. 2f (bottom) that under focusing conditions it is possible to use the DoD system to create a stable droplet train. Additionally, to maximize packing efficiency of the droplets, and thereby maximize the magnetic effect, it is important to minimize the overall size of the Ferrybot in terms of diameter. Hence choosing a core flow diameter near the diameter of the droplet is desirable. Overall, this analysis allowed us to determine the operation space for stable droplet trains aligned with sheath focusing which we then used subsequently to embed the ferrofluid droplets within the Ferrybots.

### 3.2 Ferrybot fabrication

Having established acceptable flow parameters to create the ferrofluid droplets surrounded in prepolymer solutions, the next step was the polymerization process to fix this state into a structurally-sound Ferrybot. The UV polymerization dose sets a limit on the total flow rates at which the Ferrybot can be formed. It also determines the minimum exposure needed for the ferrofluid droplets to be sealed within the Ferrybot, given the absorption/scattering of the ferrofluid particles. Lower UV polymerization doses can lead to incomplete polymerization behind the ferrofluid droplet and an open cavity in the polymerized Ferrybot resulting in the leakage of ferrofluid over time.

Our system consisted of a dual-sided collimated UV exposure setup with a 3 mm exposure area surrounding the capillary (Fig. 3a). For an exposure time (t) and a UV intensity (I_0_) at a given distance (z) from the UV source, the total dose D is given by:

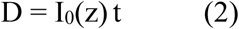

**Fig. 3.**
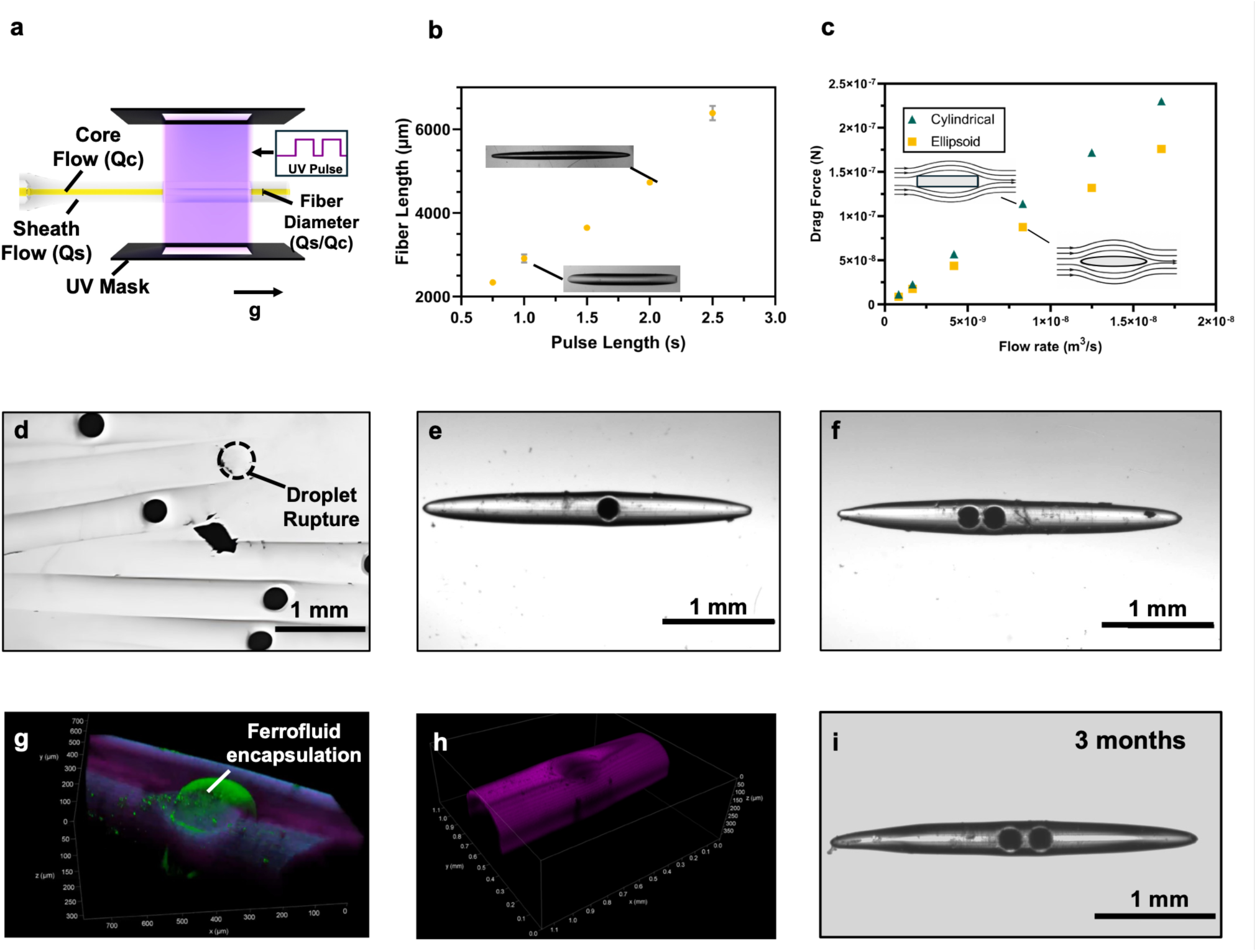
Ferrybot fabrication. (a)Layout of the Ferrybot fabrication setup (b) Ferrybot lengths as a function of UV exposure time. (c) Simulation of drag reduction for resulting oblong Ferrybots compared to similar cylindrical Ferrybots. (d) Batch of fabricated Ferrybots containing single magnetic ferrofluid core. Incomplete polymerization leads to leakage and rupture of the ferrofluid core. (e)-(f) Single and dual encapsulated magnetic cores showing programmability of Ferrybots. (g)-(h) Confocal images of Ferrybot showing the full encapsulation of the ferrofluidic core in the Ferrybot. (i) Dual core Ferrybot stored in ethanol for 3 months showing intact ferrofluid cores.

Since the polymerization time t for any given point is determined by the traversal distance within the exposure area, we can say that for a traversal distance d and flow velocity Q/A where A is the cross-sectional area and Q is the volumetric flow rate, the total dose D is given as:

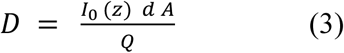

As the flow velocity increases, the total dose decreases since each point is in the exposure area for less time. For flow testing, a conservative total flow rate of 50 µL min⁻¹ was chosen to allow for sufficient traversal time to form the Ferrybots. We expected the produced Ferrybots to have an oblong tapered shape given the gradual ramp in and out of the UV exposure area as the Ferrybot travels, with the polymerized area increasing exponentially with exposure time. Based on the focusing conditions identified in Section 3.1, we used a flow rate ratio of 2.33, splitting into 15 µL min⁻¹ for the core flow and 35 µL min⁻¹ for the sheath flow. In Fig. 3b, the overall Ferrybot length increased linearly with UV pulse duration, with a minimum exposure time of 1 s required for structurally intact Ferrybots. Ferrybot length was highly reproducible, with CVs of less than 5%. This demonstrated the capability of the system to repeatedly produce Ferrybots in a consistent manner, addressing reproducibility needed for manufacturing of microbots repeatably and potentially scalably. Interestingly, laminar-flow simulations predicted lower drag forces for the oblong Ferrybot shape than for a cylindrical rod of similar dimensions (Fig. 3c). This streamlined nature of the Ferrybot could prove beneficial in navigating vasculatures as it would be less affected by the drag forces of blood flow opposing the movement of microbots^43^.

Having established the conditions that defined the outer Ferrybot structure, we next integrated programmable ferrofluid loading with the photopolymerization process. Single- and double-droplet configurations were generated (Fig. 3e–f), followed by a controlled delay that positioned the ferrofluid droplets within the UV exposure region before polymerization. This synchronization enabled reproducible encapsulation of either one or two ferrofluid droplets while maintaining similar overall Ferrybot morphology. The integrated process achieved a minimum fabrication cycle time of 15 s. Occasional fabrication failures resulted from incomplete polymerization around the ferrofluid droplets, leading to ferrofluid leakage into the surrounding medium (Fig. 3d). We next evaluated the completed Ferrybots to confirm the structural encapsulation and stability of the ferrofluid cores. Confocal imaging showed that the cores were fully enclosed within the hydrogel body (Fig. 3g). Imaging near the Ferrybot surface also revealed circular depressions adjacent to the ferrofluid cores (Fig. 3h), which likely arose from local changes in UV exposure caused by the curved capillary geometry and optical interference from the ferrofluid. To assess long-term retention, Ferrybots were stored in ethanol under gentle agitation for 3 months. The ferrofluid cores remained intact with no visible leakage or loss of magnetic function during subsequent movement testing (Fig. 3i). This stability is consistent with the PEGDA 250 network having an effective pore size smaller than the ferrofluid particles.

### 3.3 Open movement

Having established reproducible fabrication of Ferrybots with defined magnetic loading, we next characterized their fundamental modes of motion in an open fluid environment. Three complementary behaviors were evaluated: translational acceleration toward a stationary magnet, rotational response to a changing magnetic-field direction, and the maximum speed at which Ferrybots could follow a translating magnet. Together, these measurements assessed how effectively the embedded ferrofluid cores converted external magnetic inputs into linear and angular motion. Comparing one- and two-core Ferrybots also allowed us to determine how programmed magnetic loading influenced actuation strength and movement performance. We first measured translational acceleration under a fixed magnetic-field gradient. One and two-core Ferrybots were positioned at a defined starting location within a fluid-filled slot, with a 200 mT permanent magnet placed 1 mm from the surface. The slot allowed us to align the bots with the field and control the starting position of the movement. By analyzing the positional data in the video recording, we were able to determine the acceleration of both bots over time as can be seen in the snapshots of Fig. 4a. We observed a gradual nonlinear increase in acceleration of the bots towards the magnet with the two-core bot exhibiting stronger accelerations. This was to be expected since the two-core bot had higher ferrofluid content. These results demonstrate that magnetic loading can be programmed through ferrofluid core number to tune the translational response of Ferrybots under a shared magnetic field.

**Fig. 4.**
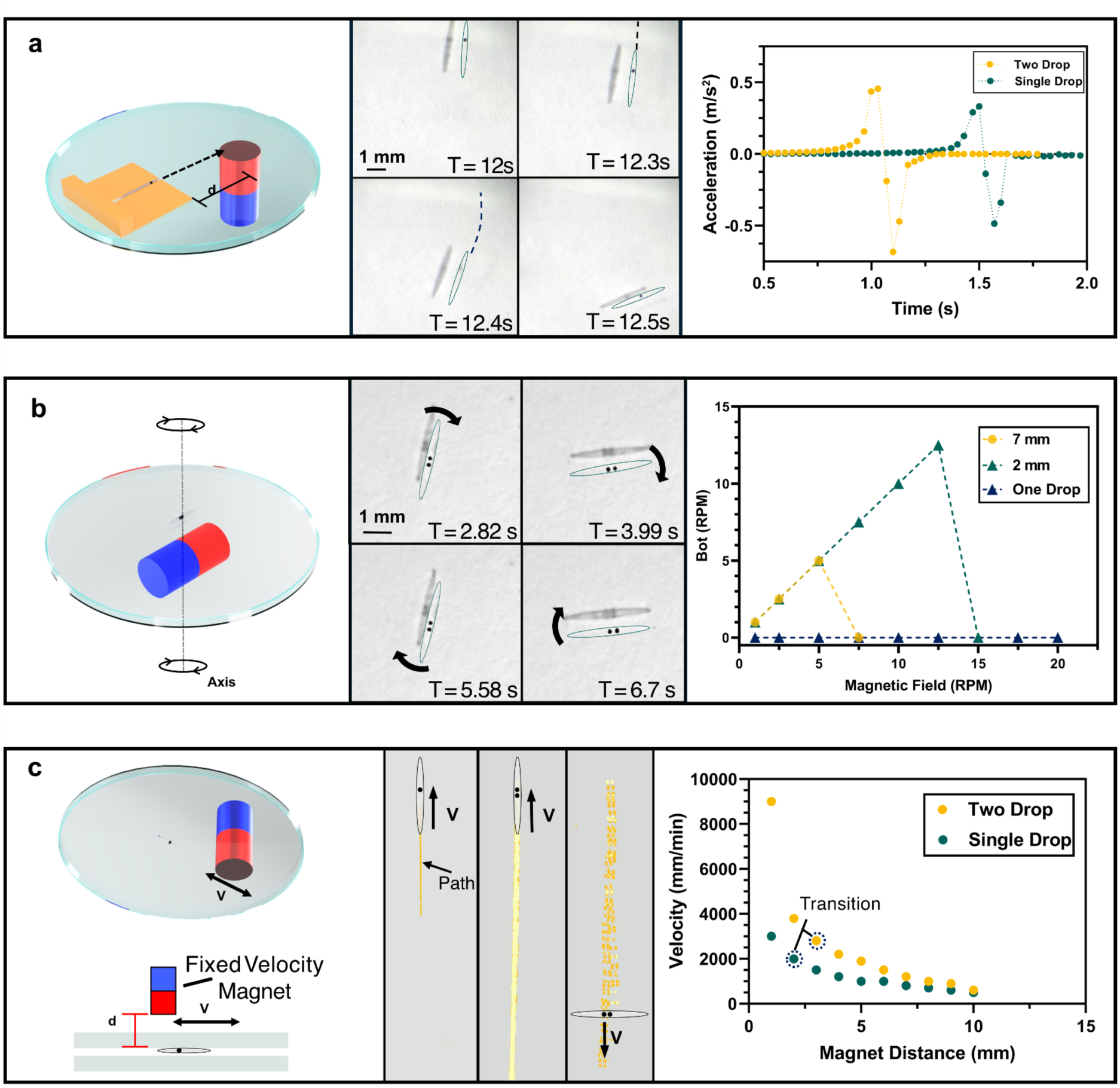
Open movement characterization. (a) Acceleration analysis for one and two-core Ferrybots using a fixed height magnet. Ferrofluid core embeddings clearly experience different force levels. (b) Rotational response of Ferrybots to increasing magnet rotation speed at different magnet distances. (c) Maximum velocity of one- and two-core Ferrybots at different magnet to Ferrybot separation distances.

We next evaluated whether Ferrybots could undergo controlled angular reorientation in response to a rotating magnetic field. A permanent magnet was mounted on a fixed-speed rotary motor beneath the sample, and its distance from the Ferrybots was varied to alter the applied field strength and gradient. Ferrybots were positioned near the magnet center, and the rotational speed was gradually increased. Under these conditions, two-core Ferrybots rotated with the magnet, whereas consistent rotation was not observed for one-core Ferrybots (Fig. 4b). With the magnet positioned 2 mm from the sample, two-core Ferrybots matched the magnet rotation rate up to 12.5 rpm (Movie S2). Above this rate, the Ferrybots could no longer maintain stable synchronous rotation. The mechanism underlying this difference was not directly measured. One plausible explanation is that the spatially separated ferrofluid cores experience unequal magnetic forces within the nonuniform field of the permanent magnet, generating a net torque about the Ferrybot center as the magnet rotates. The two-core geometry may therefore provide a larger rotational lever arm than the one-core configuration. Field-induced alignment or chaining of nanoparticles within or between the ferrofluid cores may also contribute. Further measurements or magnetic-field simulations would be required to distinguish these mechanisms.

Finally, we determined the maximum translational speed at which Ferrybots could remain coupled to and follow a moving magnetic-field source. We placed the same magnet at a fixed height as shown in Fig. 4c and moved the magnet back and forth at a specified velocity. At a certain speed, the acceleration of the bot will not be large enough to maintain the distance with the magnet nor catch up with it. We started with a separation distance of 10 mm and determined the maximum speed at which the bot could move and stay engaged throughout the process. Interestingly, both the one- and two-core bots demonstrated very fast movements, even at a 10 mm separation distance, with speeds reaching 500 mm min⁻¹ and 600 mm min⁻¹, respectively (Movie S1). At the closest separation of 1 mm both bots could move at very rapid speeds, with the one-core bot reaching 3000 mm min⁻¹ and the two-core reaching 9000 mm min⁻¹. Because 9000 mm min⁻¹ was the maximum speed of our XYZ gantry, this value represents a lower bound for two-core Ferrybots. Decreasing the separation between the magnet and Ferrybot produced a nonlinear increase in the maximum translational speed, consistent with the stronger magnetic field gradient at shorter distances. At the smallest separations, the Ferrybots also exhibited abrupt changes in orientation as they approached the edge of the cylindrical magnet. Because the field near the magnet edge is spatially nonuniform and includes substantial lateral components, the direction of the net magnetic force can shift relative to the Ferrybot’s long axis. This behavior suggests that strong local field gradients can be used not only to increase translational speed but also to induce reorientation during navigation.

### 3.4 Navigation

Having established the fundamental translational and rotational responses of Ferrybots in open fluid environments, we next evaluated whether these modes of motion could support controlled navigation through geometrically constrained settings. These experiments were designed to test path selection at channel bifurcations, maneuvering through turns, out-of-plane obstacle traversal, and selective routing of Ferrybots with different magnetic loadings. We first examined steering through a model vasculature network containing 1 mm × 1 mm channels that branched from a common inlet at angles ranging from 15° to 60° (Fig. 5a), with representative higher-angle navigation shown at 30° and 60°. We were able to steer the bot into each channel depending on the desired path (Movie S2). While preprogrammed inputs were successful for navigation, we observed that navigation through real-time control via a gaming console or through keyboard input was more consistent because it allowed the user to account for movement variations during navigation and provide appropriate feedback. Turning angles higher than 60° were typically unsuccessful due to the physical constraints of the bot’s length.

**Fig. 5.**
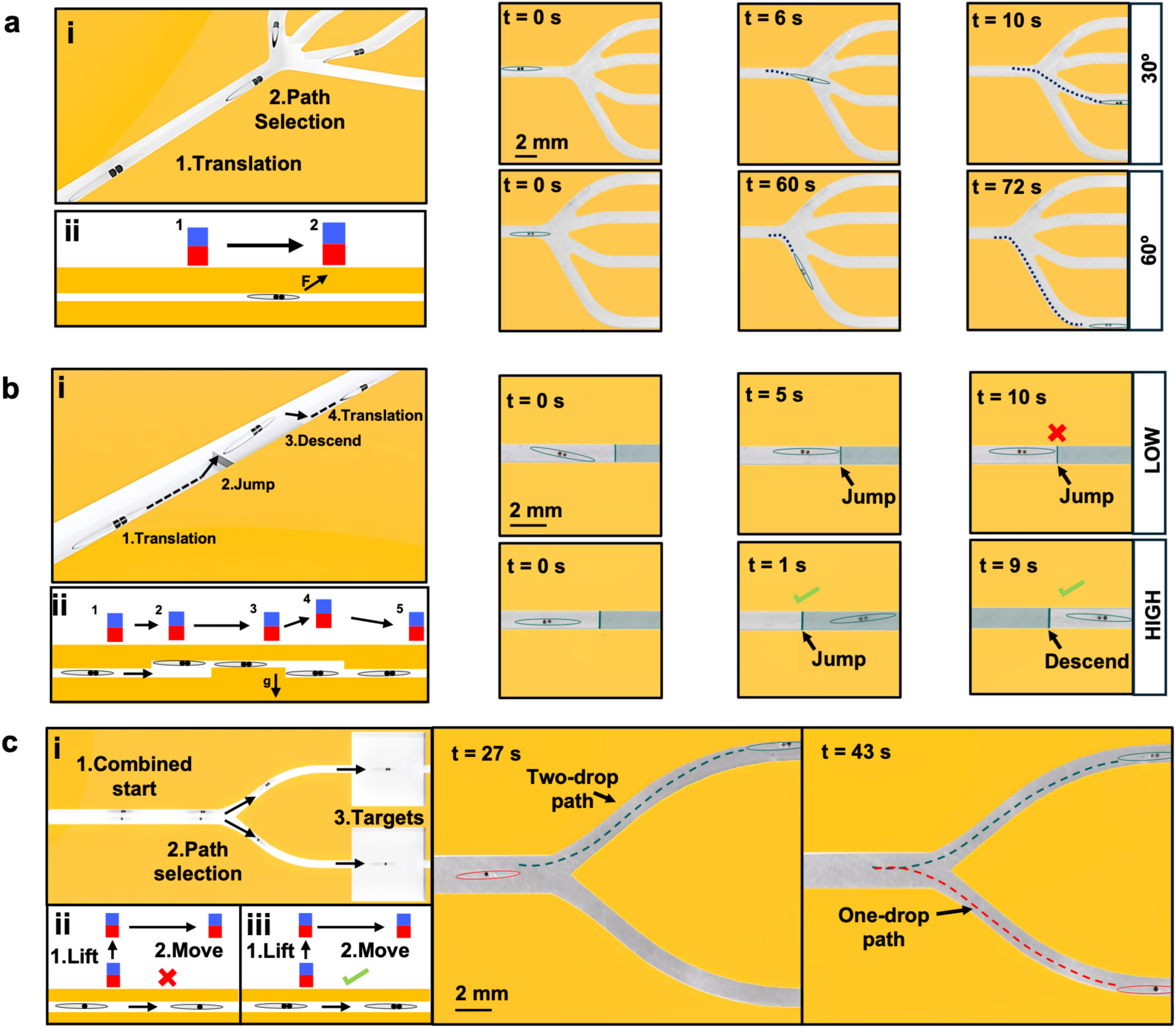
Navigation and coordination capabilities in complex 3D environments. (a) Ferrybots actuated into selected model vasculature branches ranging from 15 through 60° turns. Representative navigation through 30° and 60° branches are shown (b) Two-core Ferrybots successfully traversed out-of-plane obstacles under sufficiently strong magnetic actuation by lifting over and descending past the obstacle. (c) Demonstration of magnetically selective navigation of one and two-core Ferrybots to target different target locations in the model vasculature network.

To facilitate 3D navigation, it was important for us to address the ability for the Ferrybots to be lifted from the channel surface. We designed a channel with a hurdle (Fig. 5b) requiring vertical movement for the sake of full navigation. When the magnet was placed 5 mm from the Ferrybot (‘High field’), we could lift the bot above the hurdle after which an increase in the distance dropped the bot back on the surface (Movie S2). For distances greater than 5 mm the field was too weak to lift the bot consistently against gravity. This experiment demonstrates that the incorporated magnetic loading was sufficient to produce out-of-plane movement and obstacle traversal within the model vasculature geometry.

We next tested whether differences in magnetic core number could be used to selectively route distinct Ferrybot populations within the same channel network. One-core and two-core Ferrybots were placed together upstream of a bifurcation and assigned to separate target branches (Fig. 5c, left). With the magnet positioned farther from the channel, the more strongly responsive two-core Ferrybot engaged with the field while the one-core Ferrybot remained stationary. The two-core Ferrybot was then steered into the first target branch (Fig. 5c, center). The magnet was subsequently repositioned near the one-core Ferrybot and brought closer to increase the local field gradient, allowing the one-core Ferrybot to be engaged and directed into the second target branch (Fig. 5c, right). These results demonstrate that programmed differences in magnetic loading can enable sequential, selective navigation of heterogeneous Ferrybot populations using a shared magnetic actuation system. Such control could support deployment of Ferrybots carrying different delivery or sensing modules to multiple locations within the same environment.

### 3.5 Modular functional cargo platform

A central objective of the Ferrybot platform is to extend beyond magnetic navigation by transporting modular hydrogel cargo capable of performing application-specific functions. Rather than redesigning the magnetic vehicle for each new application, we developed a fabrication strategy that enables independently fabricated hydrogel cargo to be fused onto prefabricated Ferrybots through a common PEGDA interface. This modular approach separates microbot fabrication from cargo design, allowing functional cargo to be independently optimized before integration.

To establish this modular platform, we sought to demonstrate three key capabilities: (i) PEGDA 575 hydrogel cargo can be fused onto preformed Ferrybots, (ii) cargo integration preserves magnetic navigation, and (iii) PEGDA 575 can be independently formulated to support tunable cargo functions. As representative demonstrations of this concept, we selected programmable molecular elution and oxygen-sensing as two fundamentally different hydrogel functions to illustrate the versatility of the modular cargo architecture.

The first two capabilities were evaluated using cargo-fused Ferrybots, whereas cargo functionality was characterized using independently fabricated PEGDA 575 modules. Together, these experiments establish a modular system in which a common cargo material, such as PEGDA 575, can be independently tuned for application-specific functions and subsequently integrated with the Ferrybot without requiring redesign of the magnetic vehicle.

#### 3.5.1 Modular cargo integration

To integrate functional cargo in a modular manner, Ferrybots were positioned within a 200 µm circular PDMS mold, and a secondary PEGDA 575 hydrogel with application-specific cargo was photopolymerized directly onto one of the two tails of the Ferrybot body (Fig. 6a, b). Polymerization across the shared acrylate interface produced a continuous hydrogel structure without altering the underlying magnetic architecture of the Ferrybot. Because both the Ferrybot body and functional cargo are fabricated from PEGDA, this approach provides a straightforward and generalizable method for integrating interchangeable hydrogel modules while independently optimizing the magnetic vehicle and cargo functionality.

**Fig. 6.**
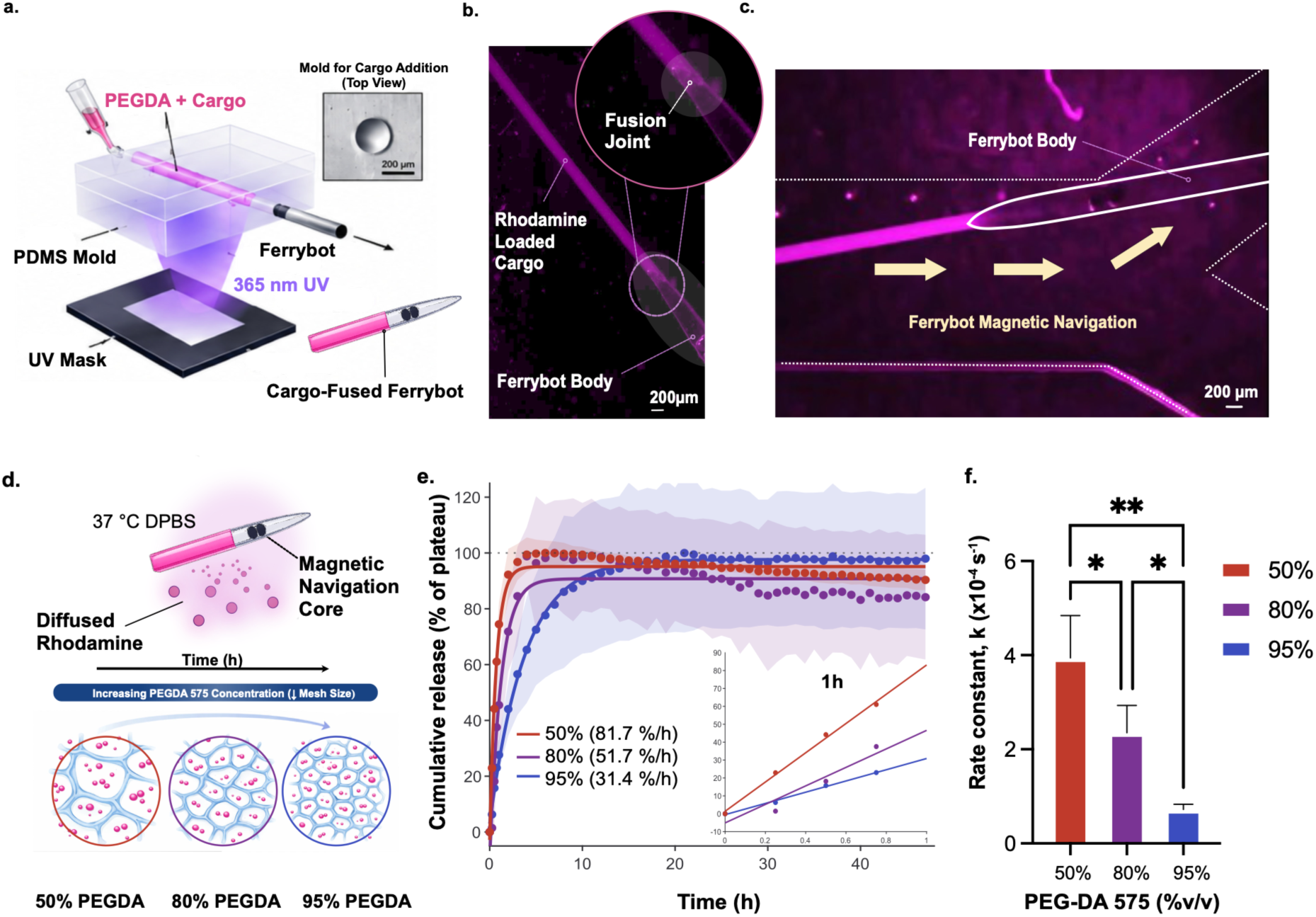
Modular functional cargo integration and programmable molecular delivery using the Ferrybot platform. (a) Soft lithographic fabrication strategy used to integrate PEGDA 575 hydrogel cargo onto Ferrybots using a circular PDMS mold. (b) Representative image of a rhodamine-loaded PEGDA 575 hydrogel cargo fused to a Ferrybot. (c) Cargo-loaded Ferrybot navigating a microfluidic channel under magnetic actuation. (d) Schematic illustrating the hypothesized relationship between PEGDA mesh size and molecular transport. (e) Molecular release profiles measured from PEGDA 575 hydrogel cargo demonstrating programmable release through hydrogel composition. (f) Apparent release constants determined from exponential fitting. Data are presented as mean ± standard deviation (SD) (n = 3). Statistical significance was determined using one-way ANOVA with Tukey’s post hoc test (*p < 0.05, **p < 0.01) via GraphPad Prism v10.

Cargo-fused Ferrybots were magnetically actuated following integration and successfully navigated microfluidic channels under external magnetic control (Fig. 6c; Movie S3), demonstrating that Ferrybots retained magnetic mobility and channel-navigation capability following cargo attachment.

#### 3.5.2 Modular molecular delivery cargo

To demonstrate that PEGDA 575 hydrogel cargo retained programmable molecular delivery functionality, rhodamine B was selected as a model small-molecule payload because its fluorescence and UV-Vis absorbance (554 nm) enable quantitative characterization of molecular transport from hydrogel matrices. These experiments were performed using independently fabricated PEGDA 575 hydrogel cargo to evaluate whether hydrogel formulation preserved predictable molecular release characteristics prior to integration. Demonstrating retained release behavior establishes the functional basis for incorporating molecular delivery modules onto the Ferrybot platform.

Previous studies have shown that increasing monomer concentration produces a denser crosslinked hydrogel network with a smaller effective mesh size, thereby restricting molecular transport and reducing solute release rates^44,45^. To investigate whether molecular release could be programmed through hydrogel formulation, rhodamine B-loaded PEGDA 575 cargo containing 50%, 80%, or 95% (v/v) PEGDA was fabricated. These formulations systematically varied hydrogel crosslink density and molecular permeability, enabling the influence of polymer composition on release kinetics to be quantified (Fig. 6d–f).

Following fabrication, rhodamine B release from PEGDA 575 hydrogel cargo was quantified over 48 h using background-corrected absorbance measurements (Fig. 6e).

All PEGDA 575 formulations exhibited exponential release characterized by an initial high rate of molecular elution followed by gradual equilibration with the surrounding medium. Release profiles were fit using a three-parameter exponential model,

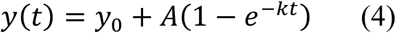

where *y*(*t*) is the measured absorbance at time *t*, *y*_0_ represents the baseline absorbance, *A* is the total absorbance change, and *k* is the apparent molecular release constant.

Molecular release was strongly dependent on PEGDA composition. The 50% PEGDA formulation exhibited the most rapid release (highest release constant, *k*), while increasing polymer concentration progressively reduced molecular transport, with the 95% PEGDA formulation exhibiting an approximately 5.5-fold lower *k*. The exponential model described the data well (R² = 0.934–0.998) across all formulations. Fits were reproducible across replicates, indicating that PEGDA composition provides a means of programming the apparent molecular release rate through hydrogel formulation (Table 1).

**Table 1.** Molecular release constants (*k*) and half-life of rhodamine B cargo over 48 h.

| **Condition** | **Release Constant, *k* (s<sup>−1</sup>)** | **Half-life, t<sub>1/2</sub> (min)** | ***R*<sup>2</sup>** |
|---|---|---|---|
| 50% PEGDA | (3.94 ± 0.90) × 10<sup>−4</sup> | 29.32 ± 6.70 | 0.973 |
| 80% PEGDA | (2.35 ± 0.58) × 10<sup>−4</sup> | 49.16 ± 12.13 | 0.934 |
| 95% PEGDA | (0.72 ± 0.11) × 10<sup>−4</sup> | 160.45 ± 24.51 | 0.998 |

##### 3.5.2.1 Early release kinetics

While the exponential model describes long-term (48 h) molecular transport, many therapeutic delivery applications are dominated by the early release period immediately following deployment. We therefore quantified the first hour of molecular release using linear regression to estimate normalized initial molecular release.

Consistent with the full release analysis, initial molecular release decreased with increasing PEGDA concentration. The 50% PEGDA formulation exhibited the highest initial release rate, whereas the 80% and 95% PEGDA formulations released molecules substantially more slowly. Coefficients of determination (R² = 0.899–0.990) indicate that early-stage molecular transport can be approximated by a constant initial rate over the first hour. Together with the exponential kinetic analysis, these findings demonstrate that PEGDA 575 hydrogel cargo retains programmable release characteristics over both early and long-term timescales, supporting its use as interchangeable molecular delivery modules for Ferrybots (Table 2).

**Table 2.** Initial molecular release analysis of rhodamine B-loaded PEGDA cargo.

| **Condition** | **1h Release Rate (%·h<sup>−1</sup>)** | ***R*<sup>2</sup>** |
|---|---|---|
| 50% PEGDA | 81.7 | 0.990 |
| 80% PEGDA | 51.7 | 0.901 |
| 95% PEGDA | 31.4 | 0.899 |

#### 3.5.3 Modular oxygen-sensing cargo

Having established that PEGDA 575 hydrogel cargo can retain programmable molecular-release behavior, we next investigated whether the same modular material architecture could support a fundamentally different function: analyte sensing. Oxygen-responsive palladium (II) meso-tetra(4-carboxyphenyl) tetrabenzoporphyrin (PdBP) was incorporated into PEGDA 575 cargo as a representative minimally invasive lifetime-based sensing cargo. Upon optical excitation at 630 nm, PdBP emits near-infrared phosphorescence at approximately 810 nm. Collisional quenching of the excited triplet state by molecular oxygen reduces the phosphorescence lifetime as oxygen concentration increases (Fig. 7a).

**Fig. 7.**
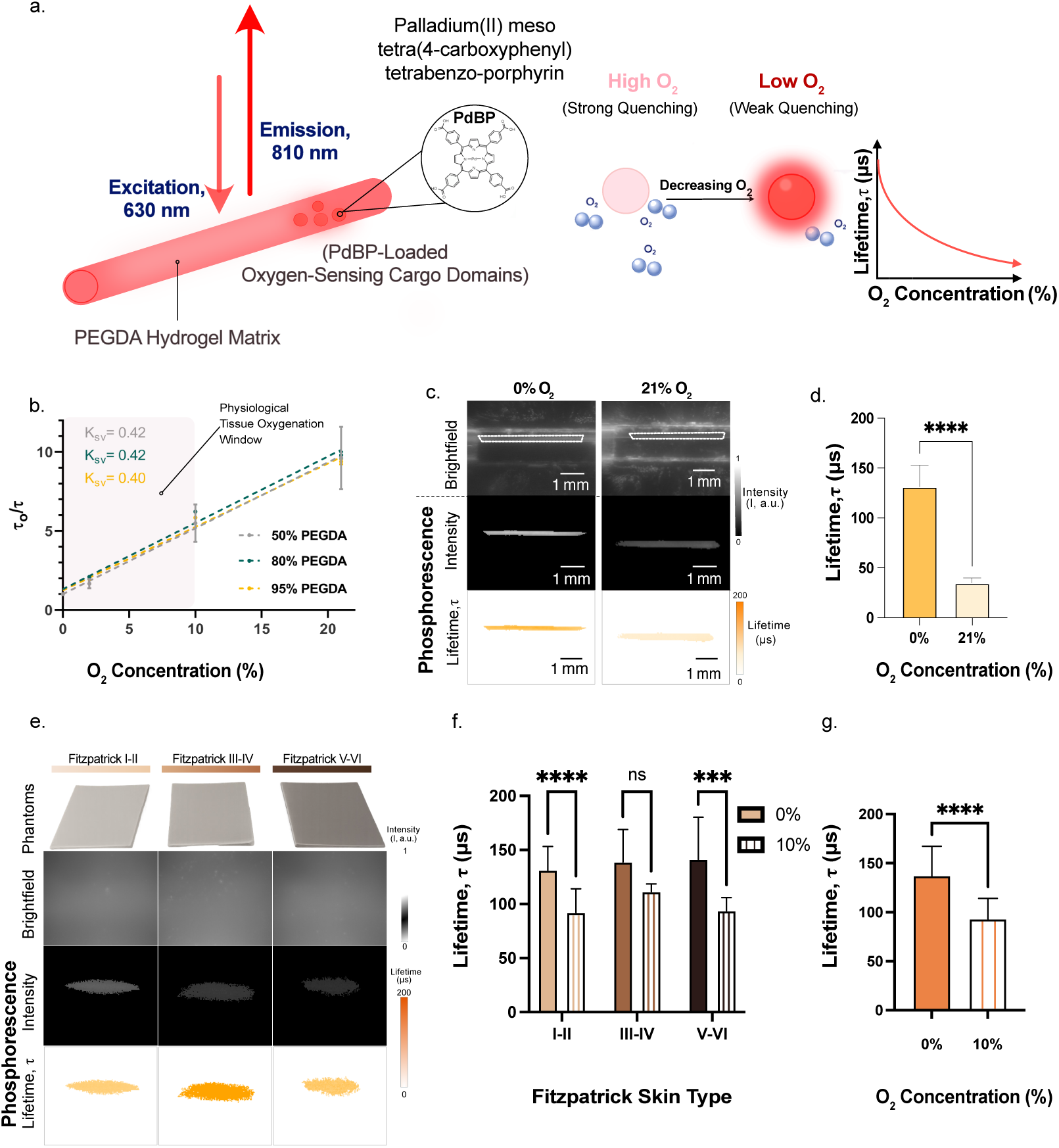
PEGDA 575 hydrogel cargo enables oxygen-responsive phosphorescence lifetime sensing across polymer formulations and through skin-mimicking phantoms. **a.** Schematic of the PdBP-loaded PEGDA 575 sensing cargo and the oxygen-dependent phosphorescence mechanism. Following excitation at 630 nm, PdBP emits near-infrared phosphorescence at approximately 810 nm. Increased oxygen concentration enhances collisional quenching of the PdBP triplet state, resulting in a shorter phosphorescence lifetime. **b.** Stern–Volmer calibration of PdBP-loaded cargos containing 50%, 80%, or 95% (v/v) PEGDA 575 over 0–21% O₂. All formulations exhibited linear oxygen-dependent responses, with *K_SV_* values of 0.42, 0.42, and 0.40 (% O₂)^-1^, respectively. The shaded region denotes the approximate physiological tissue-oxygenation range. Panel b was performed using a time resolved single pixel detector. Panels c-g were captured using a 2-D lifetime imaging camera in additional experiments. Lifetime values are compared within each experiment. **c.** Representative brightfield, phosphorescence-intensity, and fitted-lifetime images of oxygen-sensing cargo under 0% and 21% O₂ acquired using a two-dimensional lifetime imaging system. **d.** Quantification of phosphorescence lifetime under 0% and 21% O₂, demonstrating the expected reduction in lifetime with increasing oxygen concentration. **c-d.** Measurements were acquired at 25 °C (room temperature). Dashed outlines indicate the cargo boundaries. **e.** Representative skin phantoms corresponding to grouped Fitzpatrick skin types I–II, III–IV, and V–VI, together with brightfield, phosphorescence-intensity, and fitted-lifetime images of the sensing cargo acquired through each 1-mm-thick phantom (0% O_2_). Panel e represents a separate qualitative imaging experiment at 0% O₂; panels f–g summarize the subsequent measurements at 0% and 10% O₂. **f.** Phosphorescence lifetimes measured through each skin-phantom group at 0% and 10% O₂. Lifetime decreased from 0% to 10% O₂ for Fitzpatrick I–II (P < 0.0001) and V–VI (P < 0.001); III–IV did not reach significance. The O₂ response did not differ among phantom groups (O₂ skin-type interaction, P = 0.57). **g.** Aggregate lifetime response measured through the skin phantoms at 0% and 10% O₂, demonstrating retention of oxygen-dependent lifetime contrast through optically absorbing and scattering media. Scale bars, 1 mm. Statistical significance was determined using two-way ANOVA with Tukey’s post hoc test (f) and an unpaired two tailed Welch’s t-test (d, g). Statistical significance is denoted as follows: ns, not significant; \*\*\**P* < 0.001; and \*\*\*\**P* < 0.0001. Data are presented as mean ± SD. Analysis was performed via GraphPad Prism v10. Given the similar Stern–Volmer responses of the PEGDA compositions in panel (b), the 50% PEGDA 575 formulation was used for panels (c–g).

To determine whether hydrogel composition affected oxygen sensitivity, PdBP-containing cargo formulated with 50%, 80%, or 95% (v/v) PEGDA 575 was evaluated over 0–21% O₂, spanning physiologically relevant tissue oxygenation, and an air-equilibrated reference condition. Here, 21% O₂ represented the air-equilibrated condition, corresponding to approximately 257 µM dissolved oxygen under the experimental conditions. All three formulations exhibited highly linear Stern–Volmer responses, with coefficients of determination greater than 0.99 (Fig. 7b). The calculated Stern–Volmer constants were 0.42, 0.42, and 0.40 (% O₂) ⁻¹ for the 50%, 80%, and 95% PEGDA formulations, respectively.

The similar Stern–Volmer constants indicate that increasing PEGDA concentration from 50% to 95% did not measurably alter the oxygen sensitivity of the embedded PdBP probe under the conditions tested. This observation is particularly important when considered alongside the molecular-release results in Fig. 6. Whereas PEGDA concentration was used to regulate small-molecule release kinetics, oxygen sensitivity remained comparatively consistent across the same range of polymer formulations. The linear response within the approximate physiological tissue oxygenation window further supports the use of this cargo architecture for sensing oxygen levels relevant to biological applications^28,46^.

After establishing oxygen sensitivity using a time-resolved single-pixel reader, we next evaluated whether oxygen-dependent lifetime differences could be spatially visualized using a two-dimensional phosphorescence lifetime imaging system (Fig. S1)^37^. Whereas the single-pixel reader was used for quantitative Stern–Volmer calibration, the imaging system was used to spatially localize the cargo and assess the effect of skin-phantom pigmentation on signal recovery. Because the two systems differ in detection geometry and temporal resolution, absolute lifetimes and oxygen-dependent contrast in Figs. 7c–g are compared only within the imaging system and not against the calibration in Fig. 7b. PdBP-containing cargo was imaged under 0% and 21% O₂ to represent the extremes of the evaluated calibration range (Fig. 7c). The cargo remained readily identifiable in the brightfield, phosphorescence-intensity, and fitted-lifetime images. As expected, the anoxic condition produced a substantially longer lifetime than the air-equilibrated condition. Quantification confirmed a significant difference between the fitted lifetimes at 0% and 21% O₂ (Fig. 7d), demonstrating that the oxygen-responsive cargo could be spatially located while retaining its expected analyte-dependent optical response.

We subsequently investigated whether the sensing cargo could be detected through optically attenuating tissue-mimicking materials. One-millimeter-thick skin phantoms representing grouped Fitzpatrick skin types I–II, III–IV, and V–VI were positioned between the sensing cargo and the imaging system. These phantoms were selected to span increasing optical absorption and scattering conditions associated with different skin pigmentation groups; their optical properties are provided in prior works for 630 nm excitation^40^. Under 0% O₂, the cargo remained detectable through all three phantom groups, and fitted phosphorescence lifetime images, extracted from the time-resolved phosphorescence decay, could be recovered despite substantial differences in measured absolute signal intensity (Fig. 7e; Fig. S2).

The reduction in phosphorescence intensity with increasing phantom optical attenuation illustrates a key limitation of intensity-based transdermal optical measurements, which varies significantly relative to skin pigmentation (Fig. S2). In contrast, fitted lifetime response to oxygen did not depend detectably on phantom group: lifetimes at 0% O₂ were non-statistically significant across groups (131–141 µs), and two-way ANOVA showed no statistically significant main effect of phantom group (P = 0.27) or response dynamics to O_2_ % across skin phantom types (P = 0.57; Fig. 7f; Fig. S3). Because the lifetime is extracted from the shape of the phosphorescence decay rather than its amplitude, it is expected to be largely insensitive to attenuation of the collected signal.

Increasing O₂ from 0% to 10% shortened the lifetime through every phantom group (τ₀/τ = 1.25– 1.51), with significant reductions for Fitzpatrick skin types I–II (P < 0.0001) and V–VI (P < 0.001). Fitzpatrick skin types III–IV did not reach statistical significance but followed the same trend (Fig. 7f). Pooled across phantoms, lifetime decreased from 137 ± 30 µs to 93 ± 21 µs (P < 0.0001; Fig. 7g). These conditions represent an anoxic state and an oxygen level within the upper physiological tissue range, respectively, demonstrating that oxygen-dependent lifetime contrast is retained through absorbing and scattering skin phantoms irrespective of representative pigmentation.

Collectively, these findings establish PEGDA 575 as a modular cargo-loadable material capable of supporting both controlled molecular release and oxygen-responsive phosphorescence lifetime sensing in separate cargo. Although sensing and release were demonstrated in separate cargo modules, the oxygen-sensing cargo uses the same PEGDA 575 chemistry that was shown to be fusible to and navigable with the Ferrybot; these results therefore provide a material and fabrication basis for future integration of sensing and delivery cargo simultaneously. More broadly, the preservation of oxygen sensitivity across polymer formulations supports a modular design strategy in which hydrogel composition can be selected to tune cargo-specific functions without requiring redesign of the magnetic Ferrybot body.

## 4 Discussion

Magnetic microrobots are being developed for targeted delivery, sensing, manipulation, and minimally invasive intervention, but combining these functions in a practical platform remains challenging. Many reported systems are fabricated as integrated structures in which magnetic particles, structural materials, therapeutic agents, sensors, and release mechanisms must be optimized together^27^. This integration can produce sophisticated task-specific robots, but it also couples locomotion to payload composition and may require substantial redesign for each new application. Recent modular microrobot approaches have begun to address this limitation by separating a reusable actuation base from application-specific end effectors or functional components^47–49^.

Here, we developed Ferrybots as a complementary modular strategy based on compartmentalized magnetic loading and independently formulated hydrogel cargo. Rather than dispersing magnetic nanoparticles throughout the robot body, the DoD flow-lithography process encapsulates discrete ferrofluid droplets as high dose magnetic cores within a PEGDA hydrogel structure. This arrangement enables magnetic loading to be programmed through core number and spacing while preserving a surrounding hydrogel body suitable for subsequent cargo attachment. The automated process generated one- and two-core Ferrybots with a minimum cycle time of 15 s. The completed robots exhibited magnetic-loading-dependent acceleration, angular actuation, rapid translation, steering through branched channels, out-of-plane obstacle traversal, and selective engagement of one- and two-core populations. Two-core Ferrybots followed a translating magnet at speeds up to the 9000 mm min⁻¹ limit of the actuation system, although this value should be interpreted as a lower bound on the attainable speed rather than a measured physical maximum.

A second contribution is the separation of magnetic vehicle fabrication from functional cargo design. PEGDA 575 cargo was photopolymerized onto preformed Ferrybots without eliminating their magnetic mobility or channel-navigation capability. Independently fabricated cargo formulations exhibited polymer-composition-dependent release of rhodamine B and oxygen-dependent phosphorescence lifetime responses across 0–21% O₂. Lifetime differences could also be recovered through representative 1-mm-thick skin-mimicking optical phantoms. These demonstrations do not yet establish therapeutic delivery or in vivo sensing, but they show that chemically and physically distinct functions can be developed under conditions appropriate to the cargo and subsequently integrated with a common magnetic vehicle. This design logic addresses an important need in the field: avoiding complete reformulation of the microrobot whenever a new payload, sensor, or responsive material is introduced. Hydrogels are particularly attractive for this purpose because their composition can be adjusted to control molecular transport and biological interactions^50^.

Several challenges remain before Ferrybots can be applied in complex biological environments. The current navigation studies were performed in simplified, relatively large channels without physiological flow, vessel compliance, cellular interactions, or complex 3D branching geometries representative of the microvasculature. Biomedical microrobots must ultimately contend with blood-flow forces, tissue barriers, localization, retrieval or degradation, and reliable control in dynamic three-dimensional environments. Precise navigation of magnetic microrobots in such settings remains a broader challenge across the field and increasingly motivates closed-loop imaging and adaptive control approaches^51,52^. The oil-based ferrofluid and PEGDA 250 formulation used here will also require application-specific evaluation of material compatibility, ferrofluid containment, sterilization, and biological response. Likewise, the skin-phantom experiments establish optical signal recovery only through representative minimal scattering barriers; they do not yet define achievable sensing depth or performance in heterogeneous living tissue.

The most immediate next step is therefore to preserve the platform’s modularity while improving fabrication throughput, cargo integration, and validation under application-relevant conditions. Incorporating flow switching or additional precursor streams could create multiple spatially defined cargo compartments during automated fabrication, allowing combinations such as sensing and release within a single Ferrybot. Parallelized fabrication, in which quantitative measurements of production yield, core-placement accuracy, and robot-to-robot variability, will be needed to establish scalability beyond the demonstrated 15-s cycle. Navigation should be evaluated under controlled flow, in smaller and more tortuous channels, and ultimately using image-guided feedback. For oxygen sensing, photon detection depth through the skin is limited by skin tissue light absorption and scattering as well as the internal noise of the detector^53–55^. NIR-II red-shifted probes (1000–1700 nm) and improved lifetime-imaging instrumentation could increase interrogation depth, while comparison against independent oxygen measurements would establish accuracy in tissue-like environments^56–58^. Radiopaque or otherwise imageable cargo could provide complementary localization during deeper interventions.

## 5 Conclusion

Overall, Ferrybots establish a programmable magnetic hydrogel microrobot architecture in which magnetic loading, vehicle structure, and functional cargo can be designed as partially independent modules. The present study demonstrates the foundational manufacturing, locomotion, selective-navigation, cargo-integration, molecular-release, and optical-sensing capabilities of this approach. More broadly, the work suggests a route from individually optimized microrobots toward adaptable vehicle platforms that can be configured with different cargo modules for localized delivery, environmental sensing, or ultimately combined diagnostic and therapeutic functions.

## Supporting information

Schematic S1

## 6 Acknowledgements

We thank Dr. Haley L. Marks for help with confocal laser scanning microscopy at the Advanced Light Microscopy/Spectroscopy Laboratory and Leica Microsystems Center of Excellence at the California NanoSystems Institute at UCLA (RRID:SCR_022789) with funding support from NIH Shared Instrumentation Grant S10OD025017, Ajmal Ajmal and Dr. Andres Rodriguez for skin phantom fabrication at the Ramella-Roman Lab of Florida International University, NSF Major Research Instrumentation grant CHE-0722519, and NSF PATHS-UP EEC-1648451.

## 7 Supplemental Information

**Figure S1:**
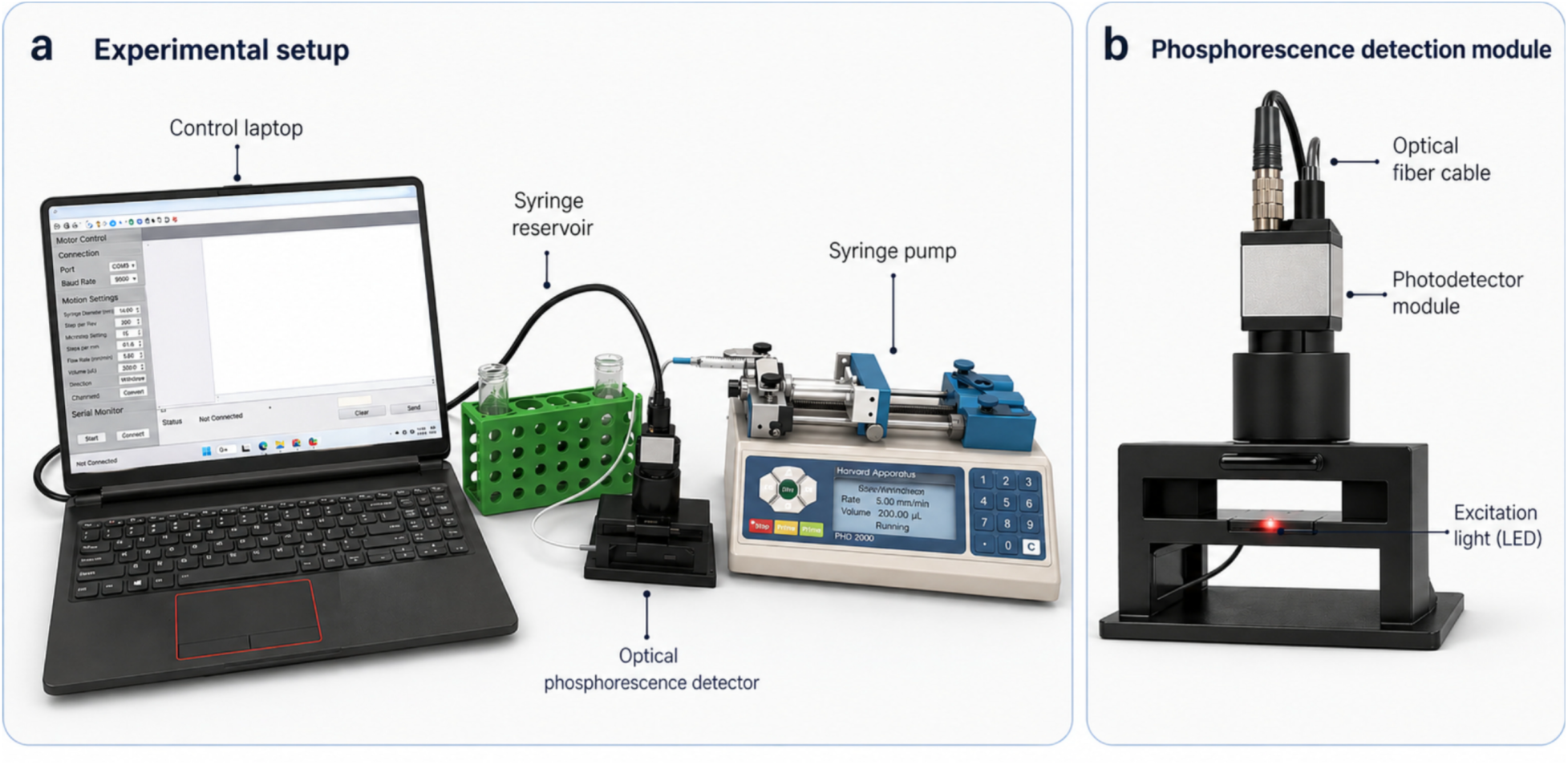
2D imaging optical setup for hydrogel cargo testing. (a) Experimental setup used for characterization of the Ferrybot oxygen-sensing platform. The system consisted of a syringe-driven flow cell integrated with a custom optical phosphorescence detection module and controlled through a laptop interface, enabling real-time imaging and analysis of oxygen-responsive sensing cargo. (b) Custom phosphorescence detection module used for two-dimensional imaging of oxygen-sensing cargo. The detector utilized 630 nm excitation and 810 nm phosphorescence emission collection to generate spatially resolved phosphorescence intensity and lifetime maps of the oxygen-sensing cargo within the sample flow cell. Images are schematic representations derived from photographs of the experimental setup, with background removed.

**Figure S2:**
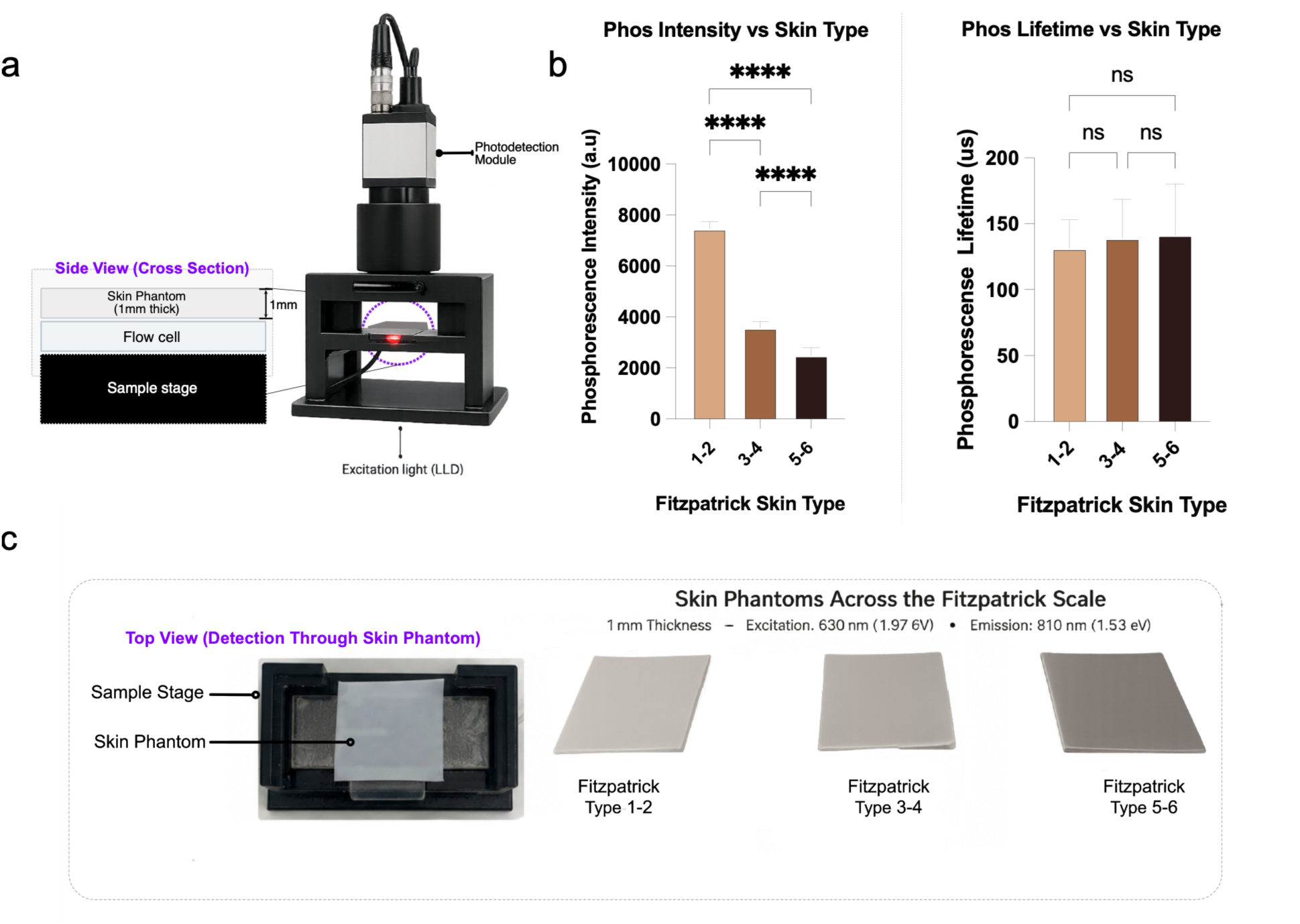
Phosphorescence intensity versus lifetime through skin phantoms. (a) Overview of the optical imaging configuration used to evaluate oxygen-sensing cargo compartments through skin mimicking phantoms. (c) Phosphorescent sensing cargo positioned within a flow cell were imaged through 1 mm-thick Fitzpatrick skin phantoms spanning skin types I–VI to assess the feasibility of non-invasive optical readout following deployment to target environments. Panels show representative top-view photographs of the imaging setup and representative skin phantom materials. (b) Phosphorescence measurements from oxygen-responsive PEGDA sensing cargo were collected through tissue-mimicking phantoms grouped by Fitzpatrick skin types I–II, III–IV, and V–VI under 0% O₂ conditions. Statistically, Phosphorescence intensity decreased significantly with increasing phantom pigmentation, whereas phosphorescence lifetime did not significantly differ across the three phantom groups. Data are presented as mean ± SD. ****p < 0.0001; ns, not significant.

**Figure S3:**
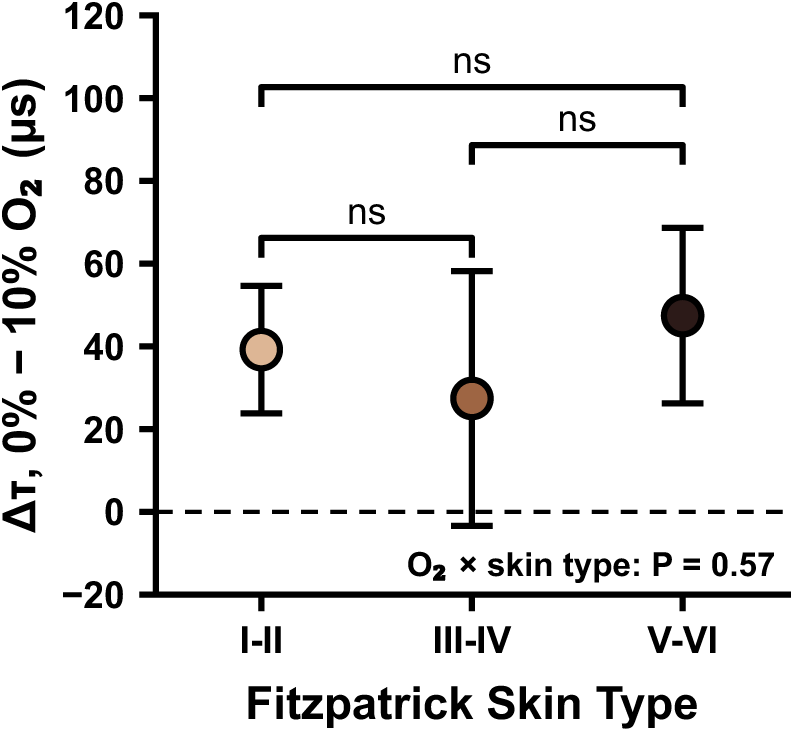
Oxygen response through each skin-phantom group. Change in phosphorescence lifetime between 0% and 10% O₂ (Δτ = τ₀% − τ₁₀%). Points show mean Δτ ± 95% confidence interval analyzed via two-way ANOVA test. The dashed line marks Δτ = 0. Δτ was 39.2 µs (95% CI 23.8–54.6) for Fitzpatrick I–II, 27.4 µs (−3.4 to 58.2) for III– IV, and 47.4 µs (26.2–68.6) for V–VI. The O₂ response did not differ significantly among phantom groups.

## Notes

### Competing Interest Statement

The authors have declared no competing interest.

